# A general mathematical framework for understanding the behavior of heterogeneous stem cell regeneration

**DOI:** 10.1101/592139

**Authors:** Jinzhi Lei

## Abstract

Stem cell heterogeneity is essential for the homeostasis in tissue development. This paper established a general formulation for understanding the dynamics of stem cell regeneration with cell heterogeneity and random transitions of epigenetic states. The model generalizes the classical G0 cell cycle model, and incorporates the epigenetic states of stem cells that are represented by a continuous multidimensional variable and the kinetic rates of cell behaviors, including proliferation, differentiation, and apoptosis, that are dependent on their epigenetic states. Moreover, the random transition of epigenetic states is represented by an inheritance probability that can be described as a conditional beta distribution. This model can be extended to investigate gene mutation-induced tumor development. The proposed formula is a generalized formula that helps us to understand various dynamic processes of stem cell regeneration, including tissue development, degeneration, and abnormal growth.

## 1 Introduction

Stem cell regeneration is an essential biological process in most self-renewing tissues during development and the maintenance of tissue homeostasis. Stem cells multiply by cell division, during which DNA is replicated and assigned to the two daughter cells along with the inheritance of epigenetic information and the partition of molecules. Unlike the accumulated process of DNA replication, inherited epigenetic information is often subjected to random perturbations; for example, the reconstruction of histone modifications and DNA methylation are intrinsically random processes of writing and erasing the modified markers [71, 91]. The stochastic inheritance of epigenetic changes during cell division can lead to stem cell heterogeneity which is important for the dynamic equilibrium of various phenotypic cells during tissue development. Accumulation of undesirable epigenetic changes may result in promoting or causing diseases [17, 18, 36, 43, 59, 60, 70, 72, 81, 92].

The heterogeneity of stem cells has been highlighted in recent years due to new technologies with single-cell resolution, which have led to the discovery of new cell types and changes in the understanding of differentiation landscapes [6, 9, 29, 50, 51, 69]. In early embryonic development, heterogeneous expression and histone modifications are correlated with correlated with cell fate and the dynamic equilibrium of pluripotent stem cells [33, 34, 68, 84]. Chromatin modifications in the human primary hematopoietic stem cell/progenitor cell (HSC/HPC) stage can lead to the dynamic equilibrium of heterogeneous and interconvertible HSCs [10, 89], as well as gene expression changes during differentiations [11]. Moreover, applications of single-cell RNA sequencing have revealed the continuous spectrum of differentiation in zebrafish [54], mice [65], and human HSCs [85]. These findings have challenged the demarcation between stem cells and progenitor cells and have led to the evolving understanding of the complex hematopoietic differentiation landscape [47, 66].

Heterogeneity plays an important role in the development of drug resistance. Cancer development is driven by evolutionary selection on somatic genetic alterations and epigenetic alterations, which result in the multistage tumorigenesis and heterogenous cancer cell phenotypes [22, 23, 39, 51, 53, 64, 90]. Tumors with different subtypes often differ in the treatment response and patient survival [13, 23, 73], and treatment stress can also induce cancer cell plasticity and drug resistance [24, 48, 57, 80, 82]. Cell plasticity is often associated with epigenetic modifications, and targeting the epigenetic regulators, such as the polycomb group protein EZH2, has been an attractive strategy in cancer treatment [20, 77, 83]. To better understand the progress of tumorigenesis and drug resistance, we need to develop predictive models of the evolutionary dynamics of cancer [3, 31].

Despite the central role of stem cell regeneration in tissue development, a quantitative investigation of the process is well beyond the ability of current technologies. Furthermore, in many fields of biological science, mathematical modeling tools have aided in improving the understanding of the principles of related processes [2, 45, 61, 62]. In 2007, Weinberg posed the following question [86]: can algebraic formulae tell us more than reasoning about the behavior of complex biological systems? Various computational models have been established in studies of tissue development and cancer systems biology under different circumstances [3, 4, 16, 25, 26, 27, 87, 88]. Nevertheless, a unified formulation that bypasses detailed assumptions is required to provide more basic logic of the biological behaviors of these complex systems. In this study, based on the general process of the cell cycle and heterogeneous stem cell regeneration, we established a general mathematical framework to formulate the dynamics of heterogeneous stem cell regeneration. The model framework includes essential cellular behaviors, including proliferation, apoptosis and differentiation/senescence; however, it bypasses the biological details of signaling pathways. The heterogeneity of stem cells and epigenetic inheritance during the cell cycle are key points in model development. Various formulas can be applied to different processes, such as embryonic development, tissue disease and degeneration, and tumor development.

The aim of this paper was to introduce a new general formula for the dynamics of stem cell regeneration with an emphasis on the effects of cell heterogeneity; therefore, a discussion of concrete conclusions based on the formula was not included. The simulation results below were included to demonstrate the potent application of the model and were not related to any actual biological processes.

## 2 Results

### 2.1 The G0 cell cycle model for homogeneous stem cell regeneration

A classical model that is used to describe the dynamics of stem cell regeneration is the G0 cell cycle model proposed in the 1970s [7, 55]. In this model, homogeneous stem cell cycles are classified into resting (G0) or proliferating (G1, S, and G2 phases and mitosis) phases (Figure 1A). During each cell cycle, a cell in the proliferating phase either undergoes apoptosis or divides into two daughter cells; however, a cell in the resting phase either irreversibly differentiates into a terminally differentiated cell or returns to the proliferating phase. This can be modeled by an age-structure model for cell numbers in the resting phase and proliferating phase. Integrating the age-structure model through the characteristic line method provides the following delay differential equation (Material and methods)

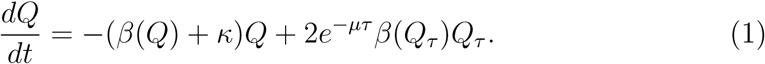

**Figure 1:**
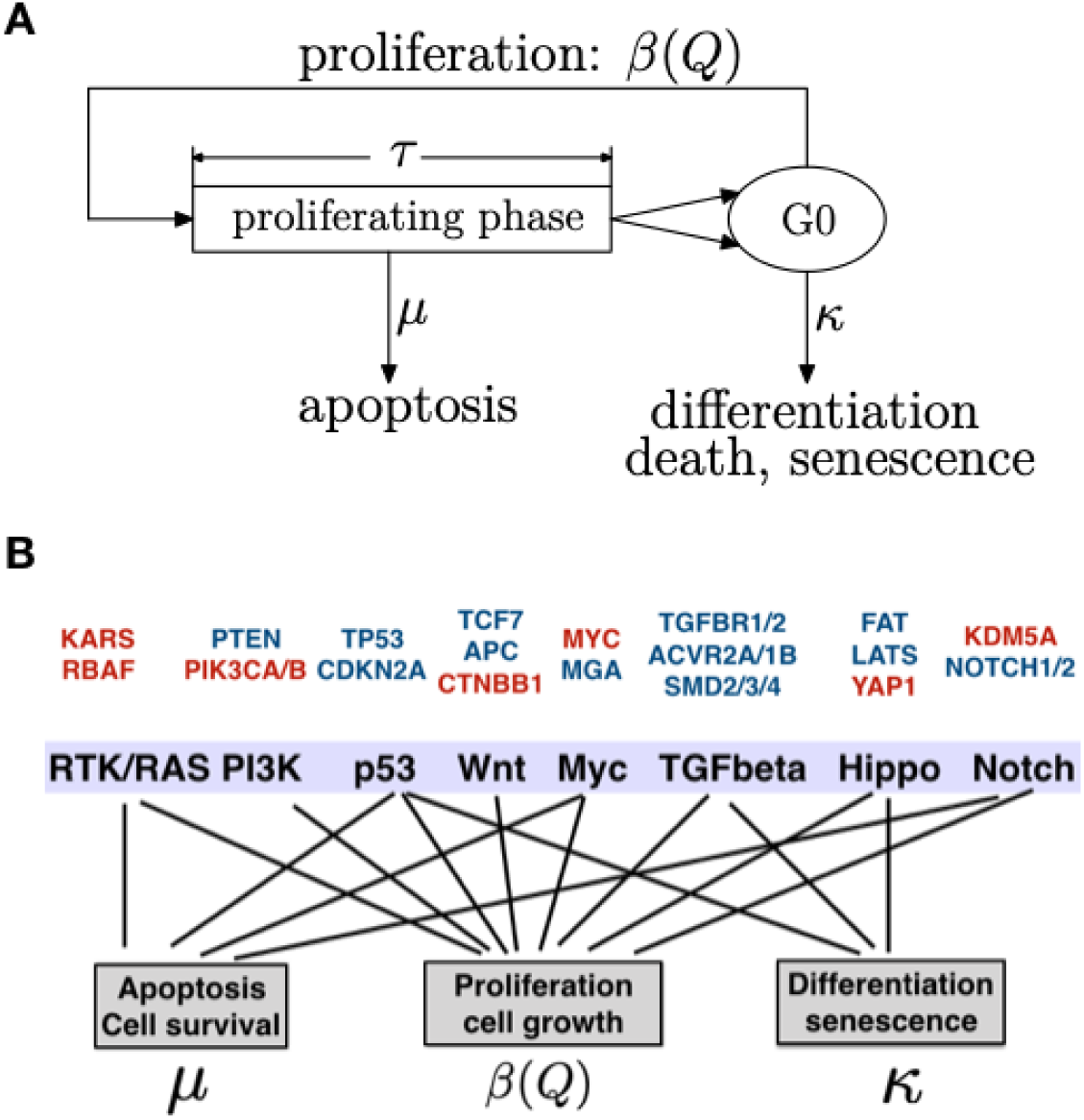
G0 cell cycle model for homogeneous stem cell regeneration. **(A)**. A schematic of the G0 model of stem cell regeneration. During stem cell regeneration, cells in the resting phase either enter the proliferation phase with the rate *β* or are removed from the resting pool with the rate *κ* due to differentiation, aging, or death. Proliferating cells undergo apoptosis with the probability *µ*. **(B)**. Oncogenic signaling pathways and their associated cell behavior and parameters in the G0 model. For each pathway, the genes are highly altered (according to the dataset in the TCGA PanCancer Atlas) by oncogenic activation (red) and tumor suppressor inactivators (blue). Details of the pathways and genes have been described by [74].

Here, *β*(*Q*) is the proliferation rate, *µ* is the apoptosis rate of cells in the proliferating phase, *τ* is the duration of the proliferating phase, and *κ* is the rate of removing cells out of the resting phase, which includes terminally differentiation, cell death, and senescence (hereafter, we call *κ* the differentiation rate for simplicity). Hereafter, the subscript indicates a time delay, *i.e., Q*_*τ*_ indicates *Q*(*t − τ*). The proliferation rate *β*(*Q*) describes how cells regulate the self-renewal of stem cells through secreted cytokines and is often given by a decrease function and *β*_0_ < *β*(*Q*) < *β*_*∞*_ (Material and method). Typically, for normal individuals, we usually have *β*_*∞*_ = lim_*Q→*+*∞*_ *β*(*Q*) = 0 because of the inhibition of the cell cycle pathway.

The G0 cell cycle model and its extensions are widely used to investigate hematopoietic stem cell dynamics [1, 19, 49, 56]; dysregulation of the apoptosis rate or differentiation rate of hematopoietic stem cells can result in serious periodic hematopoietic diseases [12]. Moreover, from (1), the stem cell dynamics are mainly determined by pathways related to stem cell proliferation, apoptosis, differentiation, senescence, and growth. Major oncogenic signaling pathways obtained from an integrated analysis of genetic alterations in The Cancer Genome Atlas (TCGA) [74] show direction connections to the coefficients *β*(*Q*), *µ*, and *κ* in (1) (Figure 1B) (Material and methods). Equation (1) is capable of describing the population dynamics of stem cell regeneration. Nevertheless, cell heterogeneity is not included in the model and has been highlighted in recent years for the understanding of cancer development and drug resistance in cancer therapy.

### 2.2 The general framework of heterogeneous stem cell regeneration

To extend the abovementioned G0 cell cycle model to include cell heterogeneity, we introduce a quantity **x** (scalar or vector) for the epigenetic state of a cell and denote *Q*(*t*, **x**) as the cell number at time *t* with state **x** (Figure 2A). In general, **x** can refer to the expression levels of marker genes, histone modifications in nucleosomes, or DNA methylations associated with DNA segments and can be measured by single-cell sequencing techniques. Specifically, we often refer to **x** as quantities that affect signaling pathways that control cell cycle progression, apoptosis, and cell growth, so that the coefficients *β, µ*, and *κ* and the duration of the proliferating phase *τ* in (1) are cell specific and dependent on the state **x** in the cell. Moreover, cells in the niche can interfere with stem cell self-renewal through released cytokines. Let *ξ*(**x**) denote the effective cytokine signal produced by a cell with state **x**, and 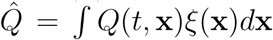 denotes the total concentration of effective cytokines that regulate cell proliferation. The proliferation rate in (1) becomes 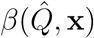.

**Figure 2:**
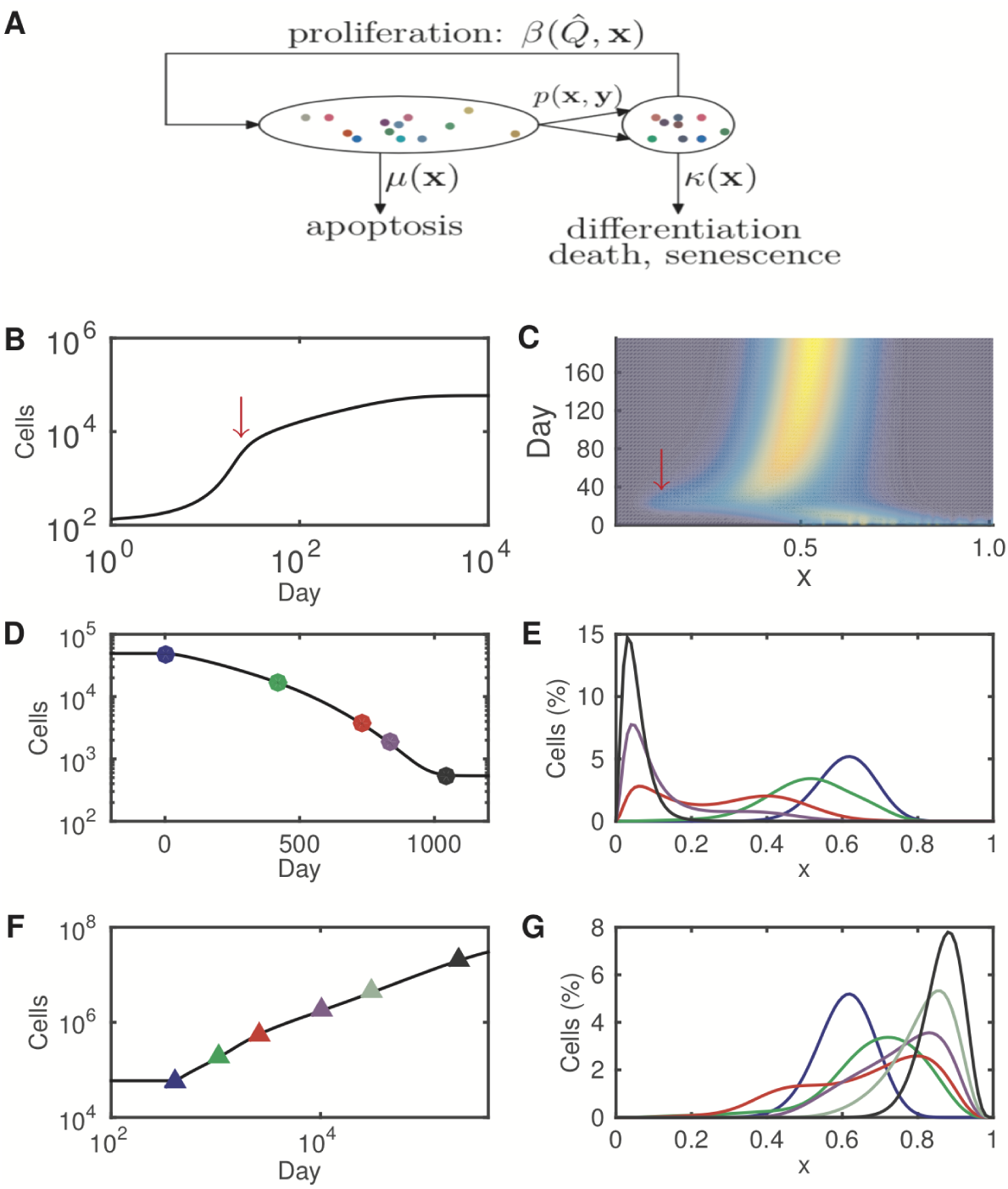
Heterogeneous stem cell regeneration. **(A)**. A schematic of the model of heterogeneous stem cell regeneration. Cell heterogeneity is represented with by the epigenetic state (**x**) indicated by dots with different colors. The dynamic rates for each cell are dependent on the epigenetic state, which varies according to cell division with the inheritance probability *p*(**x, y**). **(B)**. Simulated population dynamics for the growth process with increasing of cell numbers toward a steady state. **(C)**. The simulated evolution of cell numbers with varying heterogeneity in the growth process. The red arrows in (B) and (C) indicate a temporary increase in cell subpopulation with a low level of x at the early stage. **(D)**. Simulated population dynamics of the degeneration process with the decreasing cell numbers. **(E)**. The percentage of cells with different epigenetic states during the degeneration process. Timepoints for each curve are indicated by dots with the same colors as those shown in (D). **(F)**. Simulated population dynamics of the abnormal growth process with the increasing of cell numbers. **(G)**. The percentage of cells with different epigenetic states during the abnormal growth process. Timepoints for each curve are indicated by triangles with the same colors as those shown in (F). See Materials and methods for the simulation details.

**Figure 3:**
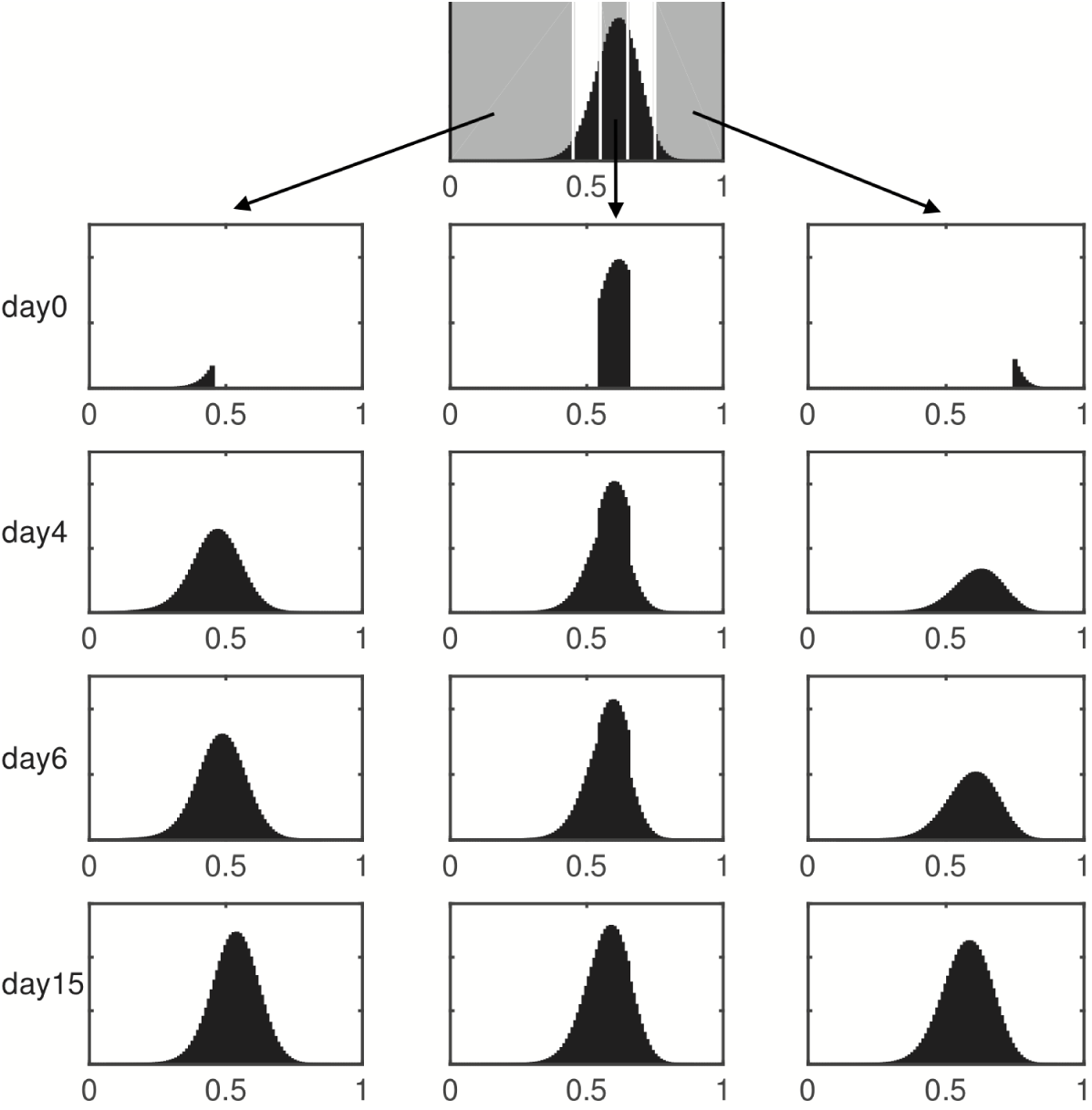
Restoration of heterogeneity from cell subpopulation fractions. Clonal cells with the highest (*x* > 0.75), middle (0.55 < *x* < 0.65), and lowest (*x* < 0.45) epigenetic states independently re-established the parental extent of clonal heterogeneity in separate simulations.

**Figure 4:**
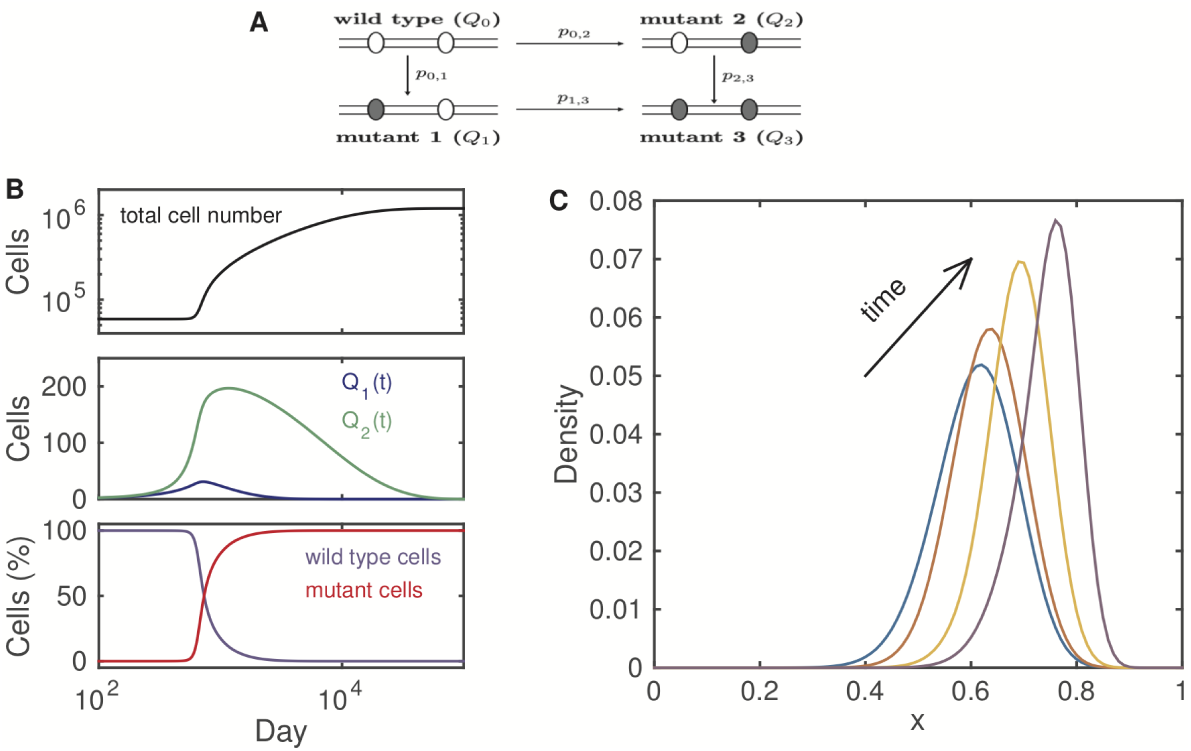
Simulated tumor development driven by mutations in proliferation and differentiation pathways. **(A)**. Cell types and mutation probability *p*_*i,j*_. Mutant 1 represents the cell type with an increased proliferation rate, mutant 2 represents the cell type with a decreasing differentiation rate, and mutant 3 represents the cell type with a double mutation. **(B)**. Evolution dynamics of total cell numbers (upper panel), mutant cells *Q*_1_(*t*) and *Q*_2_(*t*) (middle panel), and fractions of wild-type and mutant cells (lower panel). **(C)**. The evolution of cell density during tumor development.

While the cell-to-cell variability is considered, the inheritance of epigenetic states of cells during cell division is essential to shape the distribution of cell heterogeneity. Many biological processes, such as the random partition of molecules [38], random inheritance of nucleosome modification [14, 71] and DNA methylation [91, 36], can be involved in the affecting the inheritance of epigenetic states from mother to daughter cells after cell division. Many efforts have been made to model epigenetic cell memory [14, 30, 37, 38, 79]; however, it remains challenging to develop precise models for this process that is not yet clear. Nevertheless, while we overlook the biological details and focus on the changes of epigenetic states, we introduce the inheritance probability *p*(**x, y**), which represents the probability that a daughter cell of state **x** comes from a mother cell of state **y** after cell division. Therefore, *p*(**x, y**)*d***x** = 1 for any **y**. Based on the abovementioned assumptions and the similar argument to the abovementioned homogeneous model, the dynamical equation for *Q*(*t*, **x**) is as follows (Material and methods):

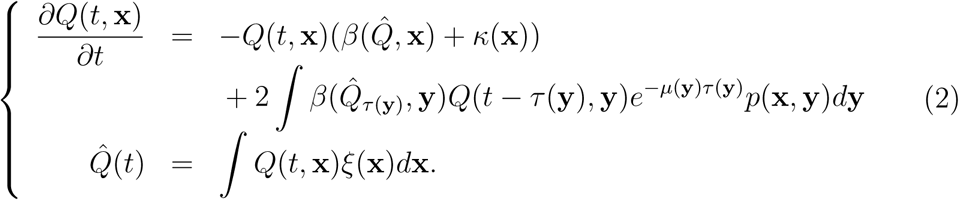

Here, the integrals are taken over all possible epigenetic states. Moreover, if we consider discrete states, such as gene mutations, we can extend the integrals to include the summation over all discrete states. Equation (2) extends the previous G0 cell cycle model and provides a general framework for heterogeneous stem cell regeneration.

Equation (2) is an autonomous system in which the rate functions and inheritance probability are not explicitly dependent on the time t. Nevertheless, while we apply the equation to situations with time dependence, such as embryo development, environmental changes, injury, and external stimuli, the time-dependent rate functions and the inheritance probability can be included in a straightforward manner. An example is shown in Figure 5 below.

**Figure 5:**
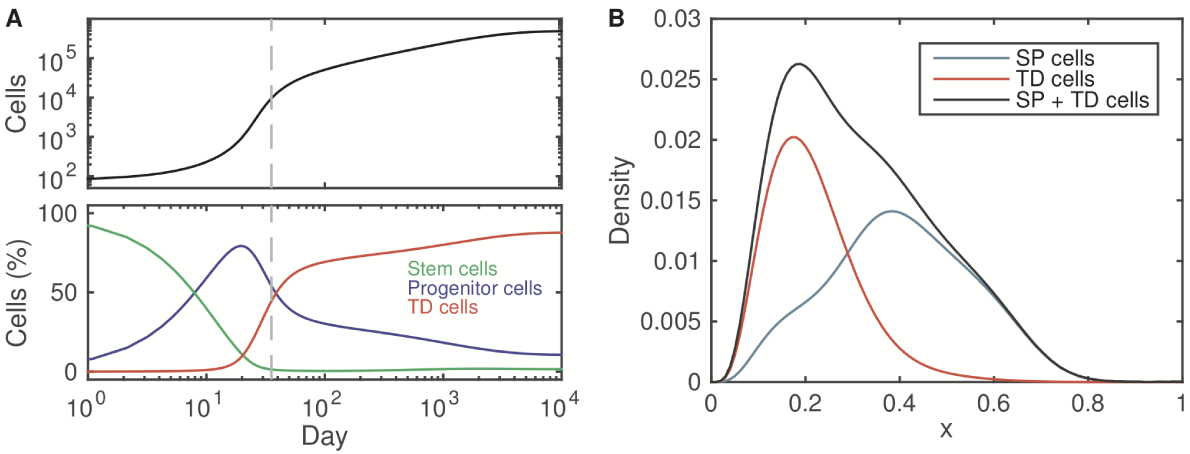
The dynamics of tissue growth. **(A).** Simulated dynamics of tissue growth beginning with 100 stem cells at day 0. Upper panel: total cell number. Lower panel: fractions of stem cells, progenitor cells, and terminally differentiated cells. **(B).** Density of the epigenetic state of three cell populations at day 35 (gray dashed line shown in (A)). Here, SP cells indicate stem and progenitor cells, and TD cells indicate terminally differentiated cells. The stem cells were defined as 0.7 ≤ *x* ≤ 1, and progenitor cells were defined as 0 < *x* < 0.7. See the Material and methods for the simulation details.

**Figure 6:**
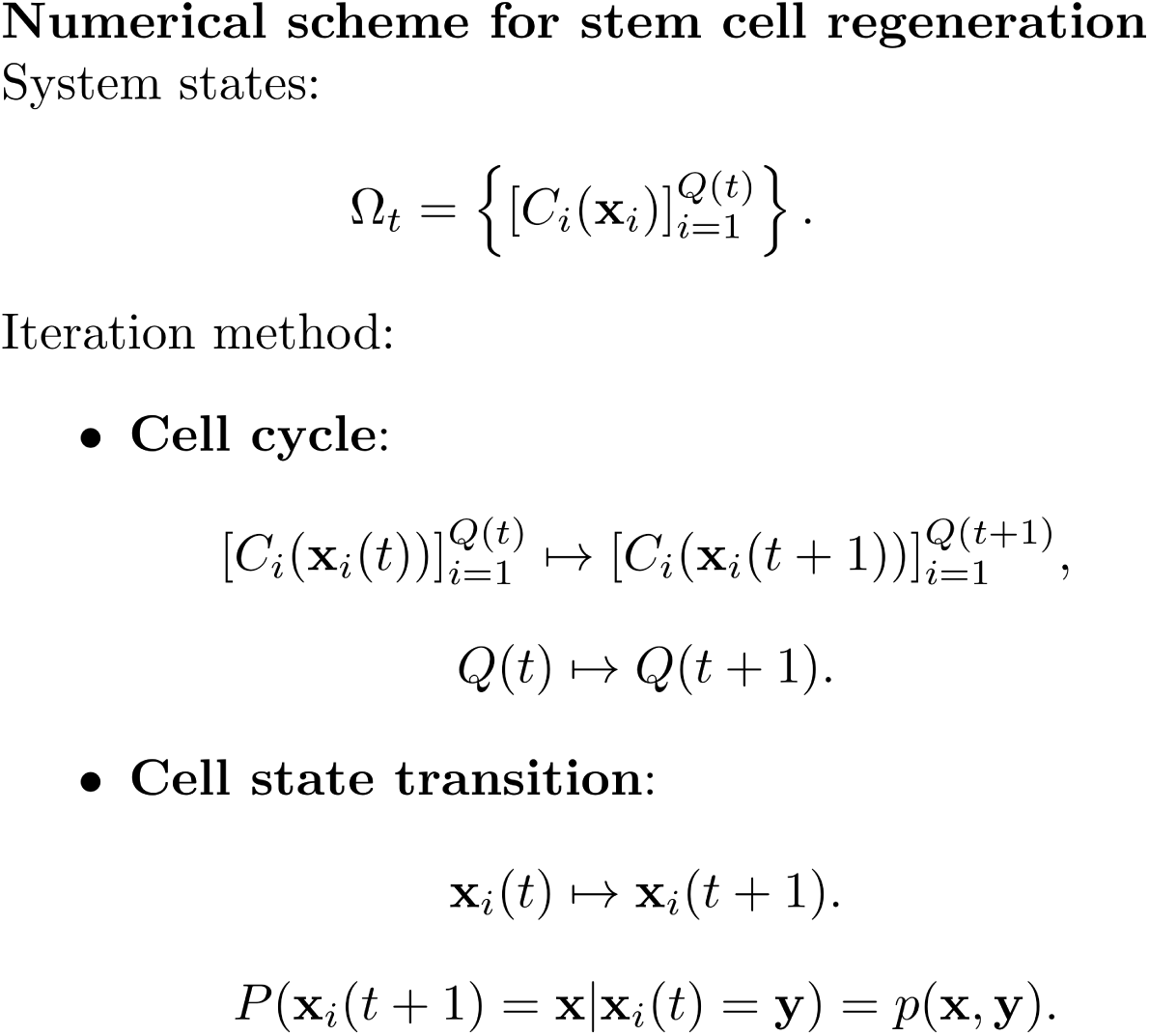
A schematic of the computational scheme for stem cell regeneration.

Based on the (2), while we introduce appropriate definitions for the dependence of kinetic rates on epigenetic states and the inheritance function, we are able to model various processes of stem cell regeneration, such as tissue growth, degeneration, and abnormal growth (Figures 2B-G). For simplicity, in all simulations shown in this paper, we consider one epigenetic state *x* (0 ≤ *x* ≤ 1) that affects only cell proliferation and differentiation in a manner similar to the stemness so that a larger value of x indicates higher stemness (Material and methods). Figure 2B-C shows the dynamics of tissue growth starting from a small population cells with high levels of stemness toward a steady state. There is a temporary transition at the early stage characterized by a rapid increase in the cell number and a subpopulation of cells with low level stemness (Figures 2B-C, red arrows). Figures 2D-E shows the dynamics of degeneration with alterations to the inheritance function, and Figures 2F-G shows the abnormal growth due to a decreasing differentiation rate and an alteration to the inheritance function. Both processes include a short-term stage of biphenotypic cell populations with both high and low stemness cells (Figure 2E and G, red curve). Moreover, the simulations show that the steady state heterogeneity can be restored from cell subpopulation fractions (Figure 3), which is in agreement with experiments that were previously explained by transcriptome-wide noise [35, 52, 89].

Equation (2) describes the evolution of the cell numbers with various epigenetic states; however, the total cell number *Q*(*t*) = ∫*Q*(*t*, **x**)*d***x** and the density of cells with different epigenetic states *f* (*t*, **x**) = *Q*(*t*, **x**)*/Q*(*t*) are relevant to the data.

From equation (2), it is easy to obtain the equations for *Q*(*t*) and *f* (*t*, **x**) as follows (see Materials and methods):

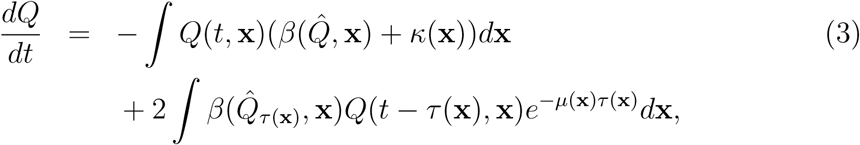

and

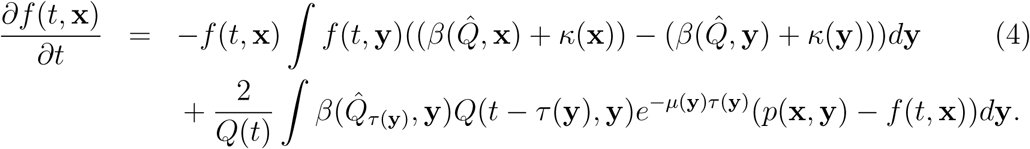

Equations (3) and (4) provide the evolution dynamics of relative cell numbers that can be obtained from experiments by single-cell sequencing or flow cytometry. Here we note that when *ξ*(**x**) = 1, we have 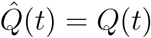, and hence, (3)-(4) provide a closed-form equation.

Here, the state variable **x** represents the epigenetic state, and *p*(**x, y**) represents the inheritance function; hence, (2) describes the dynamics with epigenetic state transitions. Nevertheless, this equation can also describe the changes in genetic alternations if we consider **x** as the genetic state and *p*(**x, y**) as the probability of point mutations. This is often the situation of genomic instability associated with cancer development [8, 32, 78, 93], and hence, the model can be used to study genetic heterogeneity in cancer development. In this paper, we focus on the equation with epigenetic state transitions and assumed that **x** always represents the epigenetic state.

### 2.3 Stochastic epigenetic state inheritance in the cell cycle

In Equations (2)-(4), the mathematical formula of epigenetic state-relevant coefficients should be expressed based on how the epigenetic states (or genes) affect the relative biological process. However, the inheritance function *p*(**x, y**) cannot be determined from the biological process of cell division. Here, we derive a phenomenological inheritance function to represent the stochastic inheritance of the epigenetic states. More specifically, let **x** = (*x*_1_, *x*_2_, …, *x*_*n*_) represent the expression level of n marker genes, and derive the inheritance function *p*_*i*_(*x*_*i*_, **y**) for each gene, and 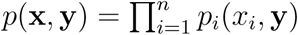.

We assume that the epigenetic state of a daughter cell is a random number with the distribution depending on that of the mother cell. In previous studies based on stochastic simulations of gene expression coupled with nucleosome modifications over multiple cell cycles [37, 40], we found that the nucleosome modification level of daughter cells, considering the nucleosome modification level of mother cells, which is normalized to the interval [0,1], can be well-described by a beta-distributed random number. Therefore, we generalized these findings and defined the inheritance function *p*_*i*_(*x*_*i*_, **y**) through the beta distribution density function as follows:

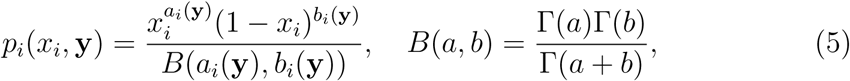

where Γ(*z*) is the gamma function and *a* and *b* are shape parameters depending on the state of the mother cell. We assumed that the mean and variance of *x*_*i*_, considering **y**, is as follows:

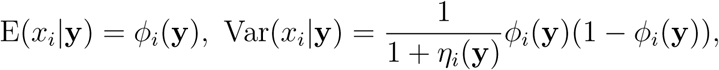

and the shape parameters are (Materials and methods)

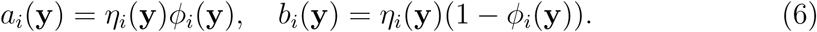

Here, we note that *ϕ*_*i*_(**y**) and *η*_*i*_(**y**) always satisfy

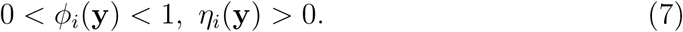

Hence, the inheritance function can be determined through the predefined functions *ϕ*_*i*_(**y**) and *η*_*i*_(**y**), often through data-driven modeling or assumptions, that satisfies (7).

### 2.4 Modeling tumor development with cell-to-cell variance

As shown in Figure 2F-G, to mimic the process of abnormal cell growth, we varied the differentiation rate and the inheritance probability. These variances to the model parameters can be a consequence of changes in the microenvironment that may affect all stem cells in the niche. Nevertheless, to model tumor development considering driver gene mutations to individual cells, we need to modify the model equations to include the mutants.

To show the framework for modeling tumor development induced by driver gene mutations, we consider the process with two types of mutations that increase the proliferation rate and decrease the differentiation rate (Figure 4A). Hence, let *Q*_*i*_(*t*, **x**) (*i* = 0, 1, 2, 3) represent the wild-type (*i* = 0) and the three mutant subpopulations (*i* = 1, 2, 3) cell counts, and *p*_*i,j*_ represents the mutation rates. For the simplicity, we assume that gene mutations occur during cell division, and two daughter cells have the same mutant type. Therefore, equation (2) can be extended as follows:

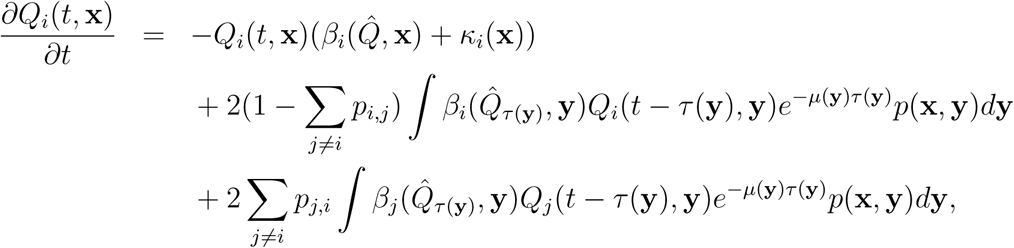

and

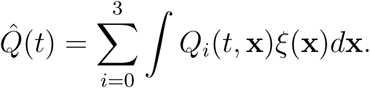

Here, we consider only the driver mutation types, and at most one mutation occurs in each cell cycle, so that only the mutation rates *p*_0,1_, *p*_0,2_ and *p*_1,3_, *p*_2,3_ are nonzero value; and otherwise, the mutation rate *p*_*i,j*_ is zero (Figure 4A).

Figure 4B-C shows the simulated dynamics. Single mutant cells occur prior to the obvious increase in the cell number, and the mutant cells eventually develop to double mutations that dominate the cell population (Figure 4B). Moreover, our simulation suggests that stemness increases with evolutionary processes when we limit the mutations to proliferation and differentiation (Figure 4C). Here, we consider only two types mutations that often occur in the precancerous stage [15, 27]. To simulate a more complicated process of cancer development, we must extend the simulation to include more mutations, such as apoptosis, DNA damage repair, and immune response pathways.

### 2.5 Modeling tissue growth with cell lineage dynamics

In the abovementioned models, we considered only cells capable of self-renewal, e.g., stem cells and progenitor cells. Nevertheless, to model tissue growth, we must include terminal differentiated cells that lose the ability to progress through the cell cycle. Therefore, let *Q*(*t*, **x**) represent cells with self-renewal ability as previously mentioned, and let *P* (*t*, **x**) represent the number of terminally differentiated cells. The terminally differentiated cells are produced from the stem and progenitor cells with the rate *κ*(**x**) and cleared with the rate *v*(**x**). Hence, equation 2 can be reformulated as follows:

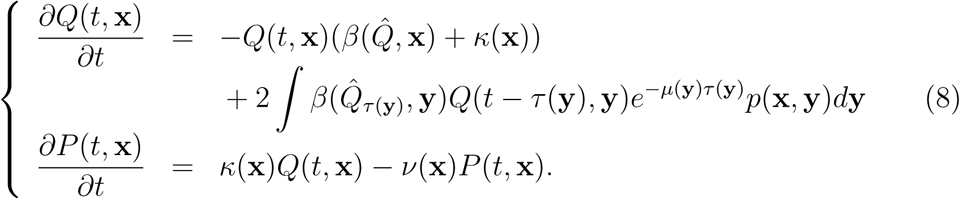

In the simulations shown in Figure 2 by considering the epigenetic state 0 ≤ *x* ≤ 1 as a stemness index and by distinguishing the stem cells from progenitor cells with the boundary *x* = *x*_0_ (Figure S1), the numbers of stem cells, progenitor cells, and terminally differentiated cells can be determined as follows:

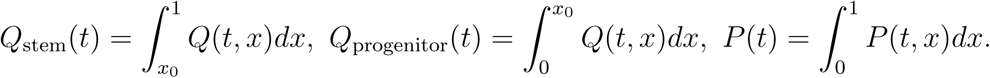

This equation provides a model of multistage cell lineages shown in previous studies [1, 21, 46]. The tissue size is given by the total cell number as follows:

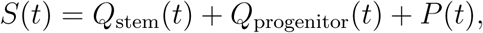

and the distribution of stemness among all tissue cells is given by

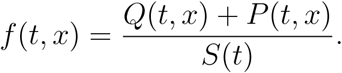

Figure 5 shows the simulated dynamics beginning with 100 stem cells, which reveal the transition between stem and progenitor cells and the differentiation to terminally differentiated cells. Figure 5B shows the density of phenotypically different cell populations among the stem cells, progenitor cells and the terminally differentiated cells.

## 3 Discussion

Stem cell and progenitor cell regeneration is a basic cellular behavior associated with development, aging, and many complex diseases in multicellular organisms. In this study, to overlook the genetic details, we established a general mathematical framework to describe the process of stem cell and progenitor cell regeneration. This framework highlights cell heterogeneity and connects heterogeneity with cellular behaviors, e.g., proliferation, apoptosis, and differentiation/senescence. Cell heterogeneity is often associated with epigenomic markers that are subject to stochastic inheritance during cell division and is described by an inheritance probability function. Hence, the framework is a multiscale model that incorporates microscopic epigenetic state and gene expressions with macroscopic tissue growth through mesoscopic cell behaviors. We believe that this formula is helpful in answering the Weinberg question [86]. Despite the generality of this formula, different assumptions regarding the kinetic rate function and the inheritance probability can be applied to describe various biological processes related to stem cell regeneration (Figure 2).

In our framework, all stem and progenitor cells are described with a single compartment model, and different phenotypic cells are not distinguished explicitly. This approach differs from differentiation tree models that are widely used to describe the maintenance of hierarchically organized tissues. Recent experimental results have challenged the discrimination between stem and progenitor cell populations and have shown a continuous spectrum of results from cell differentiation [47, 65, 85]. Stochastic state transitions between different phenotypic cells lead to a dynamic equilibrium among a population of self-renewing cells [28, 52, 75]. Our model suggested that discrimination between cell types may not be necessary to describe tissue homeostasis. Different subtypes of cells can be characterized by their kinetic rates of proliferation, apoptosis, differentiation, senescence, *etc*. For convenience, these dynamic features are referred to as the *kinotype*, which is analogous to the genotype, epigenotype, and phenotype of a cell. The kinotype of cells is often associated with specific genes enriched in the related pathways. If the relationship between kinetic rates and the expression of these genes are known, we can extend the proposed framework to include the roles of specific genes. Therefore, in the future, we aim to develop a predictable model to investigate how variations in specific genes serve to alter the long-term dynamics of tissue growth.

Although the probabilistic epigenetic inheritance was considered, equation (2) is a deterministic equation that describes the dynamics of cell densities with different epigenetic states. This model often provides information regarding the average of multiple cells. To model a single cell, we must perform stochastic simulations that explicitly account for random events. Equation (2) suggests a numerical scheme of multiscale modeling for tissue growth where a multiple cell system is represented by a collection of epigenetic states in each cell as 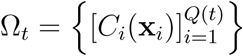. In each cell cycle, each cell undergoes proliferation, apoptosis, or terminal differentiation with a probability following the given kinetic rate so that both the system state Ω_*t*_ and the cell count *Q*(*t*) change, and the epigenetic state of each cell undergoing cell division changes according to the predefined inheritance probability *p*(**x, y**). In our previous study, this computational model was applied to model the process of inflammation-induced tumorigenesis and reproduced the two-stage tumorigenesis dynamics and revealed the competing oncogenic and onco-protective roles of inflammation. Based on the simulation results, which include the evolution of single-cell states, we were able to uncover the detailed process of cancer development.

## Material and methods

### Resource

Source MATLAB code for the study is available from https://github.com/jzlei/StemCell.

### Age-structured model and delay differential equation models

In the G0 cell cycle model, *Q*(*t*) is the number of resting-phase stem cells, *s*(*t, a*) is an age-structured quantify to represent the population of proliferating stem cells, and the age *a* = 0 is their time of entry into the proliferative state. The resting-phase cells can either reenter the proliferative phase at a rate *β*(*Q*) or differentiate into downstream cell lines at a rate *κ*. The proliferating stem cells are assumed to undergo mitosis at a fixed time *τ* after entry into the proliferating compartment and to be lost randomly at a rate *µ* during the proliferating phase. Each normal cell generates two resting-phase cells at the end of mitosis. Here, the units of cell population are often measured by the number of cells per unit body weight, *e.g.*, cells*/*kg, and the rates of proliferation, differentiation, and apoptosis are often united with day^-1^.

The above assumptions yield the following partial differential equations [49]:

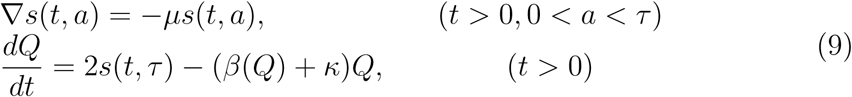

Here, ∇ = *∂/∂t* + *∂/∂a*. The boundary condition at *a* = 0 is as follows:

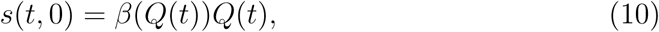

and the initial conditions are

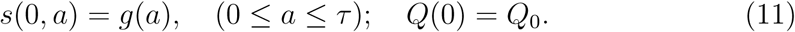

Equations (9)-(11) provide a general age-structured model of homogeneous stem cell regeneration.

By integrating (9)-(10) with the characteristic line method, we obtain the following close-form differential equation [49]:

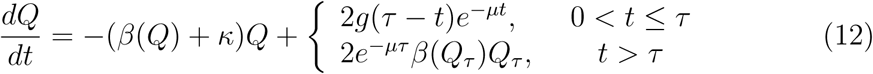

where *Q*_*τ*_ = *Q*(*t - τ*). When we consider only the long-term behavior (*t* > *τ*) and shift the original time point to *τ*, the delay differential equation model is as follows

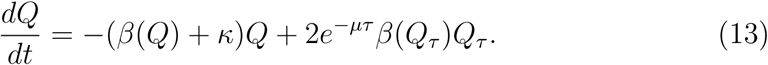

This equation describes the general population dynamics of homogeneous stem cell regeneration.

### Formulation of the proliferation rate

The effect of feedback regulation from the cell population to the proliferation rate is given by the function *β*(*Q*). Biologically, the self-renewal ability of a cell is determined by both microenvironmental conditions, *e.g.*, growth factors and various types of cytokines, and intracellular signaling pathways, e.g., growth factor receptors and cell cycle checkpoints, such as fibroblast growth factors (FGFs) and the transforming growth factor beta (TGF-*β*) family [58, 63, 67]. The exact activation pathways that regulate the self-renewal of stem cells are poorly understood. Here, we derived a phenomenological formulation based on simple but general assumptions.

There are positive and negative signals for stem cell proliferation. We assume that positive growth factors are secreted by the niche, and growth factor inhibitors are released by the cells. Different types of cytokines bind to the cell surface receptors to regulate cell behavior. Let [L] denote the concentration of ligands for growth factor inhibitor; [R], the density of free receptor; [R*\**], the density of activated receptor; *Q*, the stem cell number. The total number of receptors is

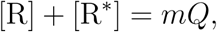

where m is the average number of receptors per cell. If n ligands are required to activate one receptor, we assume that ligands bind to the receptor following the law of mass action as follows:

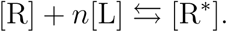

At equilibrium, we have the following equation:

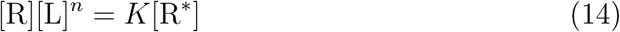

where *K* is the equilibrium constant. We assume that the activated receptors inhibit cell proliferation so that the proliferation rate is proportional to the fraction of free receptors on a cell as follows:

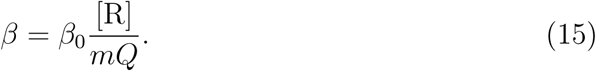

From (14)-(15), we obtain the following expression:

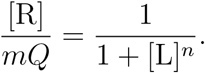

When ligands are secreted from stem cells and are cleared at a constant rate, the ligand concentration is proportional to the cell number, which gives [L] = *σQ*. Thus, we obtain the final form of the proliferation rate as follows:

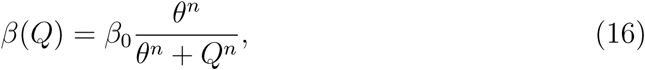

where *θ* = (1*/σ*)^1*/n*^ is the 50% effective coefficient (EC50).

From (16), the proliferation rate, which is important for tissue homeostasis, approaches 0 due to the antiproliferation signals when the cell number *Q* is sufficiently large. However, the capabilities of self-sufficiency in growth signals and insensitivity to antigrowth signals are the two characteristics of cancer that enable malignant tumor cells to escape antigrowth signals [31]. Hence, to model tumor development, the proliferation rate can be modified to include a nonzero constant *β*_1_ for self-sustained growth signals as follow:

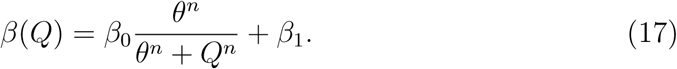

### Steady state of the G0 cell cycle model and oncogenic signaling pathways

From (13), the steady state *Q*(*t*) ≡ *Q\** is given by the equation

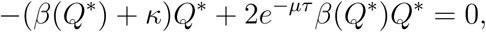

which yields either *Q\** = 0, or

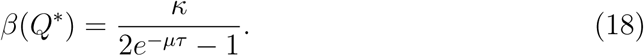

When *β*(*Q*) is given by (17), equation (13) has a unique positive steady state if and only if

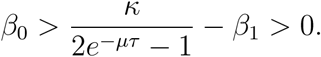

In particular, when

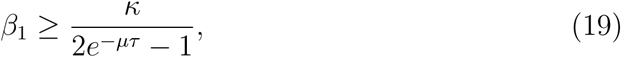

(13) has only a zero steady state, and the zero solution is unstable. Hence, all positive solutions approach infinity, which corresponds to uncontrolled growth. Therefore, the inequality (19) summarizes a general condition to have uncontrolled growth, *i.e.*, malignant tumors. Biologically, equation (19) is satisfied if there is self-sufficiency in growth signals and/or insensitivity to antigrowth signals (increasing *β*_1_), evasion of apoptosis (decreasing *µ*), and dysregulation in the differentiation and/or senescence pathways (decreasing *κ*), which are well known hallmarks of cancer [31].

### Age-structured model of heterogeneous stem cell regeneration

When heterogeneity in stem cells is considered, and assumed that the apoptosis rates of cells during cell division are dependent on the epigenetic state of the cell before entering the proliferating phase, the age-structured model (9) becomes

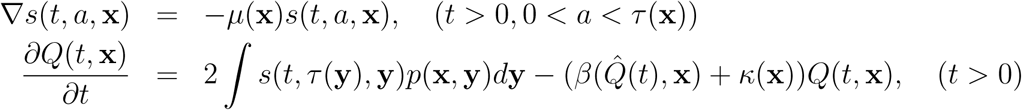

and

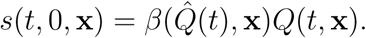

While we considered the epigenetic state **x** in the first equation as a parameter, the characteristic line method remains valid, which gives the following equation (here we show only the result of long-term behavior):

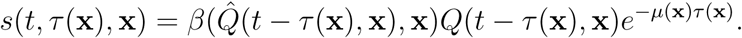

Substituting *s*(*t, τ* (**x**), **x**) into the second equation, we obtained the following equation:

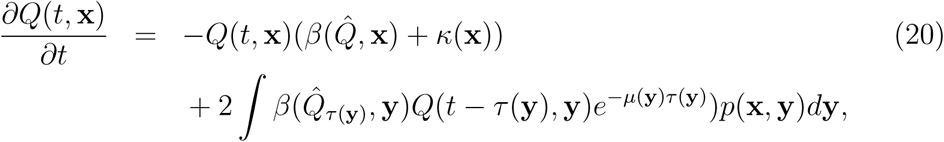

which gives the equation (2) for heterogeneous stem cell regeneration.

Let

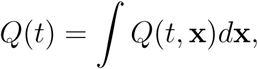

which is the total cell number, and integrate (20) with **x**. Notably, when *p*(**x, y**)*d***x** = 1, we obtain the following equation:

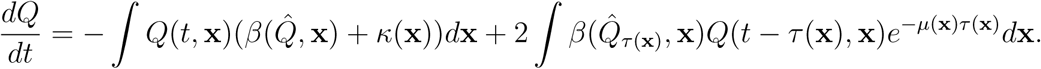

Define

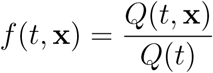

as the density of cells with a given epigenetic state **x**, then

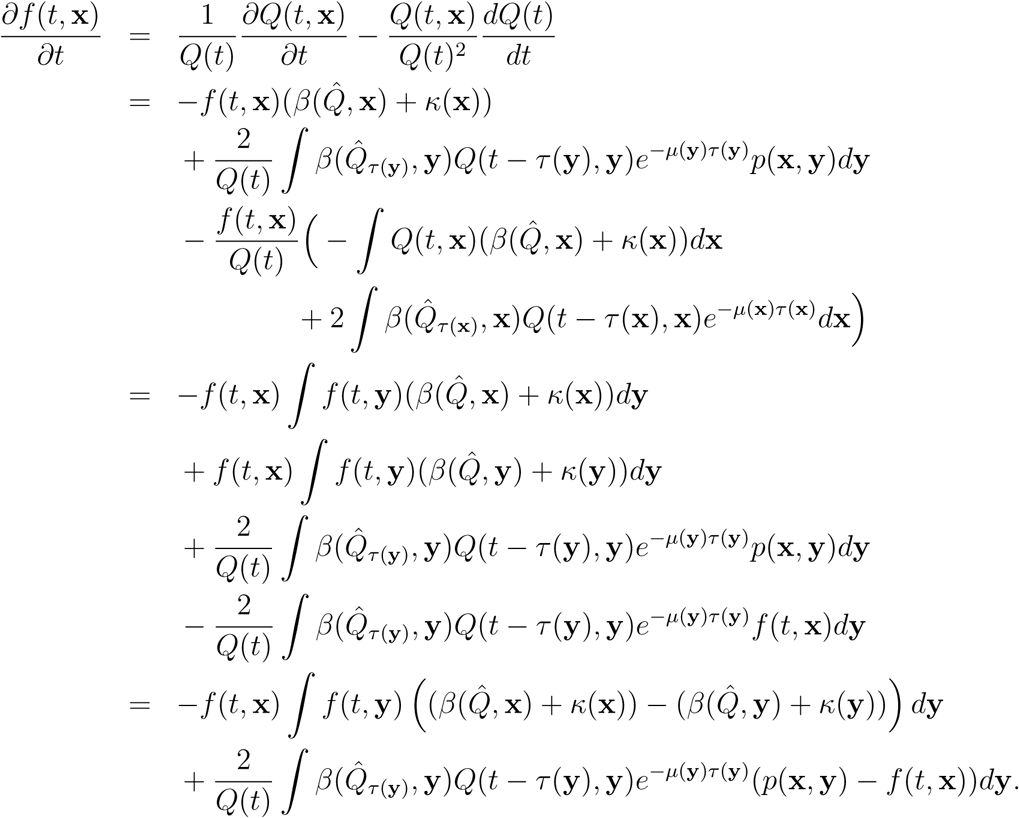

Hence, we obtained the equation

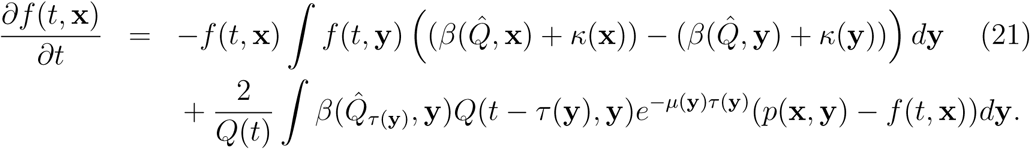

### The inheritance probability *p*(x, y)

In (2), the inheritance probability *p*(**x, y**) is essential to describe the heterogeneity of cells. The function *p*(**x, y**) is associated with the process of cell division during which the epigenetic code and molecules distribute to daughter cells through complex regulation mechanisms that are not well understood. Hence, the general mathematical formula of the function *p*(**x, y**) remains unknown. Here, we proposed an attempt to define the function based on the random inheritance of histone modifications.

In eukaryotic cells, most DNA sequences are enclosed in nucleosomes in which DNA sequences wrap around a histone octamer that is composed of one (H3-H4)2 tetramer capped by two H2A-H2B dimers. These histones can undergo diverse posttranslational covalent modifications that lead to either active or repressive gene expression activities [5, 41, 44]. The patterns of histone modification dynamically change over time, and hence define a dynamic histone code for the transcription activity. The dynamics of histone modifications consist of complex process, including nucleosome assembly, writing and erasing of the modification markers, and random inheritance during DNA replication [71, 76]. Detailed computational models for the process of histone modification and random inheritance over the cell cycle remain a challenging issue in computational biology. While we consider the main process of writing and erasing the modification markers that are modulated by the related enzymes, the kinetics of histone modification can be modeled through stochastic simulations [37, 42].

In a proposed dynamic model of histone modification [37, 42], bivalent modifications of the histone H3, the trimethylation of H3 lysine 4 (H3K4me3) and the trimethylation of H3 lysine 27 (H3K27me3), were considered. Each H3 histone can be in one of the following states: unmodified (U), modified by the activating marker H3K4me3 (A), or modified by the repressing marker H3K27me3 (R). Each nucleosome can be in one of six physically nucleosome states, which include UU, AU, UR, AA, AR, and RR. The nucleosome states dynamically change according to methylation/demethylation, which are regulated by corresponding enzymes. During DNA replication, parental histones and newly synthesized histones are randomly distributed on daughter strands. To avoid the dilution of histone markers, maintenance modifications in the new histones can be achieved by using a neighboring histone as a template [71]. Hence, writing enzyme activities are dependent on the states of neighboring nucleosomes. Thus, changes in the nucleosome state over the cell cycle due to the random distribution of histone markers during DNA replication and kinetic methylation/demethylation can be tracked with a stochastic simulation [37].

Based on the abovementioned model simulation, we are able to study how the nucleosome state of daughter cells depends on that of the mother cells. For example, considering a DNA segment with *N* nucleosomes, we counted the number (*N*_*A*_) of nucleosomes with active markers (either AA or AU) at each cycle. The simulation results suggested that considering the state of the mother cell, the active nucleosome number is a random number with a binomial distribution with the parameter (success probability *p*) dependent on the state of the mother cell as follows [37]:

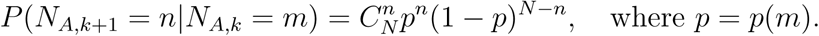

Considering the nucleosome state through the fraction (*f*_*A*_ = *N*_*A*_*/N*) of active nucleosome numbers, we can extend the binomial distribution to a continuous probability distribution defined on the interval [0, 1], which is given by the following beta distribution:

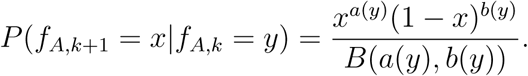

Hence, for the specific situation of the random inheritance of histone modifications during the cell cycle, we can use the beta distribution probability as the inheritance probability function *p*(*x, y*). Here, we extend this formulation to general cases and proposed (5) for the inheritance functions.

### Beta distribution

The beta distribution is a family of continuous probability distributions defined on the interval [0, 1] and is parameterized by two positive shape parameters that appear as exponents of the random variable and that control the shape of the distribution. The probability density function (PDF) of the beta distribution, for 0 ≤ *x* ≤ 1, and the shape parameters *a, b* > 0, is a power function of the variable *x* and of its reflection (1 − *x*) as follows:

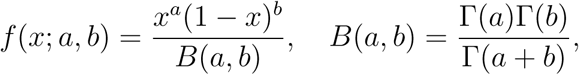

where Γ(*z*) is the gamma function.

For a random variable *X* beta-distributed with parameters *a* and *b*, which is denoted by *X* ∼ beta(*a, b*), the mean and variance are as follows:

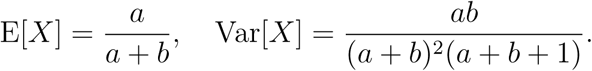

Then, it is easy to obtain

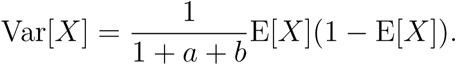

Hence, if we assume

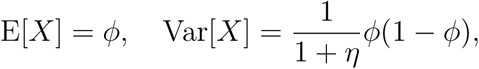

then

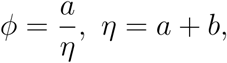

which gives

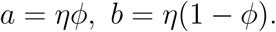

This gives equation (6) to determine the shape parameters from the functions *ϕ*_*i*_(**y**) and *η*_*i*_(**y**).

### Simulations for stem cell regeneration

Here, we present a simple example to show the numerical scheme to simulate stem cell regeneration based on the proposed model equations.

We consider a situation with one epigenetic state *x* (0 ≤ *x* ≤ 1) that affects only cell proliferation and differentiation so that only the rates *β* and *κ* are dependent on the epigenetic state *x*. Therefore, we have the following model equation:

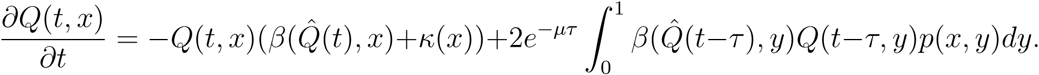

Here, *ξ*(*x*) = 1 so that

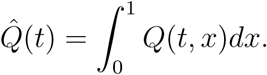

To specify the rate functions *β* and *κ*, we assume that the state *x* affects the proliferation and differentiation rates in a manner similar to the stemness so that large value of *x* indicates the stem cells with a low proliferation rate, an intermediate value *x* indicates progenitor cells with a high proliferation rate, and the terminated differentiation rate *κ* is a decreasing function of *x* that approaches zero when *x* is large. This is mathematically expressed as follows:

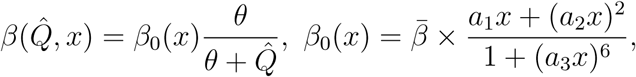

and

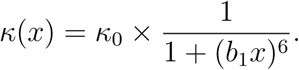

The inheritance probability function *p*(*x, y*) is defined from the beta distribution density function as with predefined function *ϕ*(*y*) and *η*(*y*) as follows:

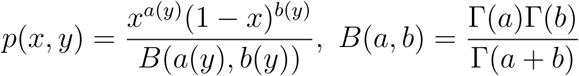

and

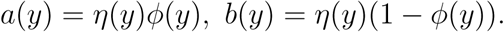

Figure 7 shows the functions *β*_0_(*x*), *κ*(*x*), and *p*(*x, y*).

**Figure 7:**
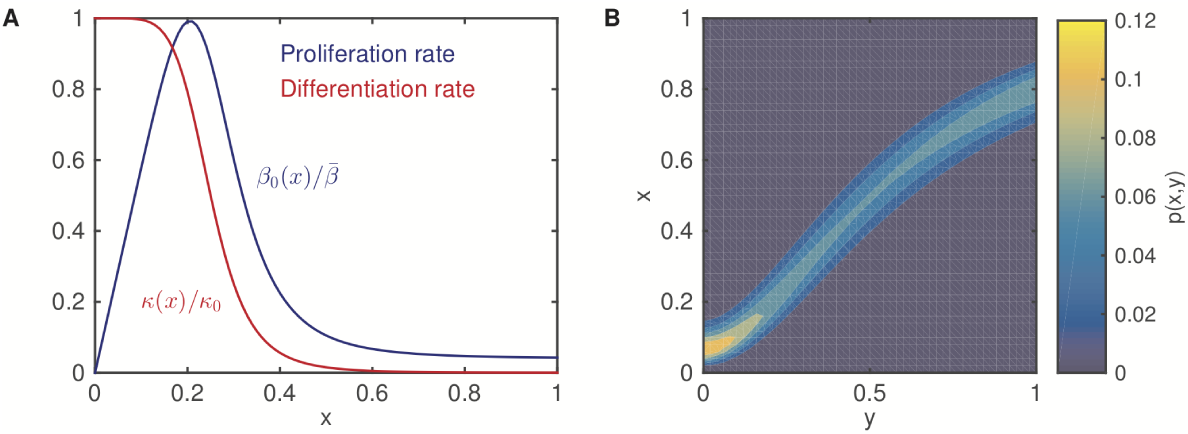
Example of the rate functions. (A). The proliferation rate *β*_0_(*x*) and the differentiation rate *κ*(*x*). (B). The inheritance probability *p*(*x, y*). Here, the parameters are *a*_1_ = 5.8, *a*_2_ = 2.2, *a*_3_ = 3.75, *b*_1_ = 4.0, and 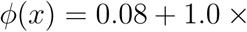 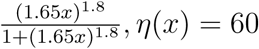.

In the simulations shown in Figure 2 the parameters are as follows: *µ* = 2.0 × 10^-4^ h^-1^, *τ* = 20 h, *θ* = 10^3^cells, *a*_1_ = 5.8, *a*_2_ = 2.2, *a*_3_ = 3.75, *b*_1_ = 4.0, *η*(*x*) = 60, and

- in (B)-(C): 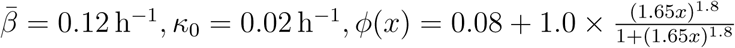;
- in (D)-(E): 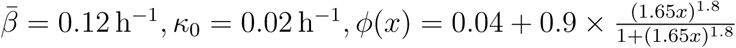;
- in (F)-(G):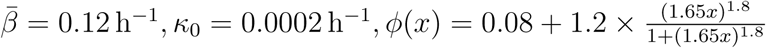.

In the simulation show in Figure 3 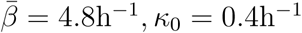, *κ*_0_ = 0.4h^-1^, and other parameters are the same as those shown in Figure 2B.

In the simulation shown in Figure 4 we set the wild-type cell parameters according to those shown in Figure 2B and *p*_0,1_ = *p*_0,2_ = 0.5 × 10^-4^, *p*_1,3_ = *p*_2,3_ = 1 × 10^-4^. For mutation 1, the proliferation rate is twice of wild-type cells, and for mutation 2, the differentiation rate is 1*/*10 of wild-type cells.

In the simulation shown in Figure 5 we set *x*_0_ = 0.7, the differentiation rate increases from 0 to normal level following 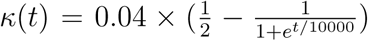, and the other parameters are the same as those shown in Figure 2B.

## Acknowledagement

This work was support by the National Natural Science Foundation of China (NSFC 91730301 and 11831015).

## References

[1] Mostafa Adimy, Oscar Angulo, Catherine Marquet, and Leila Sebaa. A mathematical model of multistage hematopoietic cell lineages. DCDS-B, 19(1):1–26, January 2014.

[2] Philipp M Altrock, Lin L Liu, and Franziska Michor. The mathematics of cancer: integrating quantitative models. Nat Rev Cancer, 15(12):730–745, November 2015.

[3] Niko Beerenwinkel, Chris D Greenman, and Jens Lagergren. Computational Cancer Biology: An Evolutionary Perspective. PLoS Comput Biol, 12(2):e1004717, February 2016.

[4] Niko Beerenwinkel, Roland F Schwarz, Moritz Gerstung, and Florian Markowetz. Cancer evolution: mathematical models and computational inference. Syst. Biol., 64(1):e1–25, January 2015.

[5] L Bintu, J Yong, Y E Antebi, K McCue, Y Kazuki, N Uno, M Oshimura, and M B Elowitz. Dynamics of epigenetic regulation at the single-cell level. Science, 351(6274):720–724, February 2016.

[6] Florian Buettner, Kedar N Natarajan, F Paolo Casale, Valentina Proserpio, Antonio Scialdone, Fabian J Theis, Sarah A Teichmann, John C Marioni, and Oliver Stegle. Computational analysis of cell-to-cell heterogeneity in single-cell RNA-sequencing data reveals hidden subpopulations of cells. Nat Biotechnol, 33(2):155–160, February 2015.

[7] F J Burns and I F Tannock. On the existence of a G0-phase in the cell cycle. Cell Proliferation, 3(4):321–334, 1970.

[8] Rebecca A Burrell, Nicholas McGranahan, Jiri Bartek, and Charles Swanton. The causes and consequences of genetic heterogeneity in cancer evolution. Nature, 501(7467):338–345, September 2013.

[9] Andrew Butler, Paul Hoffman, Peter Smibert, Efthymia Papalexi, and Rahul Satija. Integrating single-cell transcriptomic data across different conditions, technologies, and species. Nat Biotechnol, 36(5):411–420, April 2018.

[10] Hannah H Chang, Martin Hemberg, Mauricio Barahona, Donald E Ingber, and Sui Huang. Transcriptome-wide noise controls lineage choice in mammalian progenitor cells. Nature, 453(7194):544–547, May 2008.

[11] Kairong Cui, Chongzhi Zang, Tae-Young Roh, Dustin E Schones, Richard W Childs, Weiqun Peng, and Keji Zhao. Chromatin signatures in multipotent human hematopoietic stem cells indicate the fate of bivalent genes during differentiation. Stem Cell, 4(1):80–93, January 2009.

[12] David C Dale and Michael C Mackey. Understanding, treating and avoiding hematological disease: better medicine through mathematics? Bull Math Biol, 77(5):739–757, April 2015.

[13] Felipe De Sousa E Melo, Xin Wang, Marnix Jansen, Evelyn Fessler, Anne Trinh, Laura P M H de Rooij, Joan H de Jong, Onno J de Boer, Ronald van Leersum, Maarten F Bijlsma, Hans Rodermond, Maartje van der Heijden, Carel J M van Noesel, Jurriaan B Tuynman, Evelien Dekker, Florian Markowetz, Jan Paul Medema, and Louis Vermeulen. Poor-prognosis colon cancer is defined by a molecularly distinct subtype and develops from serrated precursor lesions. Nat Med, 19(5):614–618, May 2013.

[14] Ian B Dodd, Mille A Micheelsen, Kim Sneppen, and Geneviéve Thon. Theoretical analysis of epigenetic cell memory by nucleosome modification. Cell, 129(4):813–822, May 2007.

[15] Jarno Drost, Richard H van Jaarsveld, Bas Ponsioen, Cheryl Zimberlin, Ruben van Boxtel, Arjan Buijs, Norman Sachs, Rene M Overmeer, G Johan Offerhaus, Harry Begthel, Jeroen Korving, Marc van de Wetering, Gerald Schwank, Meike Logtenberg, Edwin Cuppen, Hugo J Snippert, Jan Paul Medema, Geert J P L Kops, and Hans Clevers. Sequential cancer mutations in cultured human intestinal stem cells. Nature, 521(7550):43–U329, 2015.

[16] Huijing Du, Yangyang Wang, Daniel Haensel, Briana Lee, Xing Dai, and Qing Nie. Multiscale modeling of layer formation in epidermis. PLoS Comput Biol, 14(2):e1006006, February 2018.

[17] Matthias Farlik, Florian Halbritter, Fabian Müller, Fizzah A Choudry, Peter Ebert, Johanna Klughammer, Samantha Farrow, Antonella Santoro, Valerio Ciaurro, Anthony Mathur, Rakesh Uppal, Hendrik G Stunnenberg, Willem H Ouwehand, Elisa Laurenti, Thomas Lengauer, Mattia Frontini, and Christoph Bock. DNA Methylation Dynamics of Human Hematopoietic Stem Cell Differentiation. Cell stem cell, 19(6):808–822, December 2016.

[18] Adam E Field, Neil A Robertson, Tina Wang, Aaron Havas, Trey Ideker, and Peter D Adams. DNA Methylation Clocks in Aging: Categories, Causes, and Consequences. Mol Cell, 71(6):882–895, September 2018.

[19] Catherine Foley and Michael C Mackey. Dynamic hematological disease: a review. J Math Biol, 58(1-2):285–322, 2009.

[20] Joseph A Fraietta, Christopher L Nobles, Morgan A Sammons, Stefan Lundh, Shannon A Carty, Tyler J Reich, Alexandria P Cogdill, Jennifer J D Morris-sette, Jamie E DeNizio, Shantan Reddy, Young Hwang, Mercy Gohil, Irina Kulikovskaya, Farzana Nazimuddin, Minnal Gupta, Fang Chen, John K Everett, Katherine A Alexander, Enrique Lin-Shiao, Marvin H Gee, Xiaojun Liu, Regina M Young, David Ambrose, Yan Wang, Jun Xu, Martha S Jordan, Katherine T Marcucci, Bruce L Levine, K Christopher Garcia, Yangbing Zhao, Michael Kalos, David L Porter, Rahul M Kohli, Simon F Lacey, Shelley L Berger, Frederic D Bushman, Carl H June, and J Joseph Melenhorst. Disruption of TET2 promotes the therapeutic efficacy of CD19-targeted T cells. Nature, 558(7709):307–312, June 2018.

[21] Erika Gaspari, Annika Franke, Diana Robles-Diaz, Robert Zweigerdt, Ingo Roeder, Thomas Zerjatke, and Henning Kempf. Paracrine mechanisms in early differentiation of human pluripotent stem cells: Insights from a mathematical model. Stem Cell Res, 32:1–7, October 2018.

[22] M. Gerlinger, A.J. Rowan, S. Horswell, J. Larkin, D. Endesfelder, E. Gronroos, P. Martinez, N. Matthews, A. Stewart, P. Tarpey, I Varela, B Phillimore, S Begum, N.Q. McDonald, A Butler, D Jones, K Raine, C Latimer, C.R. Santos, M Nohadani, A.C. Eklund, B Spencer-Dene, G Clark, L Pickering, G Stamp, M Gore, Z Szallasi, J Downward, P.A. Futreal, and C. Swanton. Intratumor Heterogeneity and Branched Evolution Revealed by Multiregion Sequencing. N. Engl. J. Med., 367(10):976–976, September 2012.

[23] Alice Giustacchini, Supat Thongjuea, Nikolaos Barkas, Petter S Woll, Benjamin J Povinelli, Christopher A G Booth, Paul Sopp, Ruggiero Norfo, Alba Rodriguez-Meira, Neil Ashley, Lauren Jamieson, Paresh Vyas, Kristina Anderson, Åsa Segerstolpe, Hong Qian, Ulla Olsson-Strömberg, Satu Mustjoki, Rickard Sandberg, Sten Eirik W Jacobsen, and Adam J Mead. Single-cell transcriptomics uncovers distinct molecular signatures of stem cells in chronic myeloid leukemia. Nat Med, 23(6):692–702, June 2017.

[24] Thomas Graf. Differentiation plasticity of hematopoietic cells. Blood, 99(9):3089–3101, May 2002.

[25] James M Greene, Doron Levy, Sylvia P Herrada, Michael M Gottesman, and Orit Lavi. Mathematical Modeling Reveals That Changes to Local Cell Density Dynamically Modulate Baseline Variations in Cell Growth and Drug Response. Cancer Research, 76(10):2882–2890, May 2016.

[26] Philip Greulich and Benjamin D Simons. Dynamic heterogeneity as a strategy of stem cell self-renewal. Proc Natl Acad Sci USA, 113(27):7509–7514, June 2016.

[27] Yucheng Guo, Qing Nie, Adam L MacLean, Yanda Li, Jinzhi Lei, and Shao Li. Multiscale modeling of inflammation-induced tumorigenesis reveals competing oncogenic and onco-protective roles for inflammation. Cancer Research, 77(22):6429–6441, September 2017.

[28] Piyush B Gupta, Christine M Fillmore, Guozhi Jiang, Sagi D Shapira, Kai Tao, Charlotte Kuperwasser, and Eric S Lander. Stochastic State Transitions Give Rise to Phenotypic Equilibrium in Populations of Cancer Cells. Cell, 146(4):633–644, December 2010.

[29] Adam L Haber, Moshe Biton, Noga Rogel, Rebecca H Herbst, Karthik Shekhar, Christopher Smillie, Grace Burgin, Toni M Delorey, Michael R Howitt, Yarden Katz, Itay Tirosh, Semir Beyaz, Danielle Dionne, Mei Zhang, Raktima Raychowdhury, Wendy S Garrett, Orit Rozenblatt-Rosen, Hai Ning Shi, Omer Yilmaz, Ramnik J Xavier, and Aviv Regev. A single-cell survey of the small intestinal epithelium. Nature, 551(7680):333–339, November 2017.

[30] Jan O Haerter, Cecilia Lövkvist, Ian B Dodd, and Kim Sneppen. Collaboration between CpG sites is needed for stable somatic inheritance of DNA methylation states. Nucleic Acids Res, 42(4):2235–2244, January 2014.

[31] D Hanahan and R A Weinberg. The hallmarks of cancer. Cell, 100(1):57–70, January 2000.

[32] Douglas Hanahan and Robert A Weinberg. Hallmarks of cancer: the next generation. Cell, 144(5):646–674, March 2011.

[33] R David Hawkins, Gary C Hon, Chuhu Yang, Jessica E Antosiewicz-Bourget, Leonard K Lee, Que-Minh Ngo, Sarit Klugman, Keith A Ching, Lee E Edsall, Zhen Ye, Samantha Kuan, Pengzhi Yu, Hui Liu, Xinmin Zhang, Roland D Green, Victor V Lobanenkov, Ron Stewart, James A Thomson, and Bing Ren. Dynamic chromatin states in human ES cells reveal potential regulatory sequences and genes involved in pluripotency. Cell Res, 21(10):1393–1409, October 2011.

[34] Katsuhiko Hayashi, Susana M Chuva de Sousa Lopes, Fuchou Tang, and M Azim Surani. Dynamic equilibrium and heterogeneity of mouse pluripotent stem cells with distinct functional and epigenetic states. Stem Cell, 3(4):391–401, October 2008.

[35] Martin Hoffmann, Hannah H Chang, Sui Huang, Donald E Ingber, Markus Loeffler, and Joerg Galle. Noise-Driven Stem Cell and Progenitor Population Dynamics. PLoS ONE, 3(8):e2922, August 2008.

[36] Steve Horvath. DNA methylation age of human tissues and cell types. Genome Biol., 14(10):R115, 2013.

[37] Rongsheng Huang and Jinzhi Lei. Dynamics of gene expression with positive feedback to histone modifications at bivalent domains. Int. J. Mod. Phys. B, 4:1850075, November 2017.

[38] Dann Huh and Johan Paulsson. Non-genetic heterogeneity from stochastic partitioning at cell division. Nat Genet, 43(2):95–100, January 2011.

[39] Mariam Jamal-Hanjani, Gareth A Wilson, Nicholas McGranahan, Nicolai J Birkbak, Thomas B K Watkins, Selvaraju Veeriah, Seema Shafi, Diana H Johnson, Richard Mitter, Rachel Rosenthal, Max Salm, Stuart Horswell, Mickael Escudero, Nik Matthews, Andrew Rowan, Tim Chambers, David A Moore, Samra Turajlic, Hang Xu, Siow-Ming Lee, Martin D Forster, Tanya Ahmad, Crispin T Hiley, Christopher Abbosh, Mary Falzon, Elaine Borg, Teresa Marafioti, David Lawrence, Martin Hayward, Shyam Kolvekar, Nikolaos Panagiotopoulos, Sam M Janes, Ricky Thakrar, Asia Ahmed, Fiona Blackhall, Yvonne Summers, Rajesh Shah, Leena Joseph, Anne M Quinn, Phil A Crosbie, Babu Naidu, Gary Middleton, Gerald Langman, Simon Trotter, Marianne Nicolson, Hardy Remmen, Keith Kerr, Mahendran Chetty, Lesley Gomersall, Dean A Fennell, Apostolos Nakas, Sridhar Rathinam, Girija Anand, Sajid Khan, Peter Russell, Veni Ezhil, Babikir Ismail, Melanie Irvin-Sellers, Vineet Prakash, Jason F Lester, Malgorzata Kornaszewska, Richard Attanoos, Haydn Adams, Helen Davies, Stefan Dentro, Philippe Taniere, Brendan O’Sullivan, Helen L Lowe, John A Hartley, Natasha Iles, Harriet Bell, Yenting Ngai, Jacqui A Shaw, Javier Herrero, Zoltan Szallasi, Roland F Schwarz, Aengus Stewart, Sergio A Quezada, John Le Quesne, Peter Van Loo, Caroline Dive, Allan Hackshaw, Charles Swanton, and TRACERx Consortium. Tracking the Evolution of Non-Small-Cell Lung Cancer. N. Engl. J. Med., 376(22):NEJMoa1616288–2121, April 2017.

[40] Xiaopei Jiao and Jinzhi Lei. Dynamics of gene expression based on epigenetic modifications. Communications in Information and Systems, 18(3):125–148, 2018.

[41] Tony Kouzarides. Chromatin modifications and their function. Cell, 128(4):693–705, February 2007.

[42] Wai Lim Ku, Michelle Girvan, Guo-Cheng Yuan, Francesco Sorrentino, and Edward Ott. Modeling the dynamics of bivalent histone modifications. PLoS ONE, 8(11):e77944, 2013.

[43] Marta Kulis and Manel Esteller. DNA methylation and cancer. Adv. Genet., 70:27–56, 2010.

[44] Monika Lachner, Roderick J O’Sullivan, and Thomas Jenuwein. An epigenetic road map for histone lysine methylation. Journal of Cell Science, 116(Pt 11):2117–2124, May 2003.

[45] Arthur D Lander. How Cells Know Where They Are. Science, 339(6122):923–927, 2013.

[46] Arthur D Lander, Kimberly K. Gokoffski, Frederic Y M Wan, Qing Nie, and Anne L. Calof. Cell lineages and the logic of proliferative control. PLoS biology, 7(1):e15, January 2009.

[47] Elisa Laurenti and Berthold Göttgens. From haematopoietic stem cells to complex differentiation landscapes. Nature, 553(7):418–426, January 2018.

[48] Clémentine Le Magnen, Michael M Shen, and Cory Abate-Shen. Lineage Plasticity in Cancer Progression and Treatment. Annu Rev Cancer Biol, 2:271–289, March 2018.

[49] Jinzhi Lei and Michael C Mackey. Multistability in an age-structured model of hematopoiesis: Cyclical neutropenia. J Theor Biol, 270(1):143–153, February 2011.

[50] Hanna Mendes Levitin, Jinzhou Yuan, and Peter A Sims. Single-Cell Transcriptomic Analysis of Tumor Heterogeneity. Trends in Cancer, 4(4):264–268, April 2018.

[51] Huipeng Li, Elise T Courtois, Debarka Sengupta, Yuliana Tan, Kok Hao Chen, Jolene Jie Lin Goh, Say Li Kong, Clarinda Chua, Lim Kiat Hon, Wah Siew Tan, Mark Wong, Paul Jongjoon Choi, Lawrence J K Wee, Axel M Hillmer, Iain Beehuat Tan, Paul Robson, and Shyam Prabhakar. Reference component analysis of single-cell transcriptomes elucidates cellular heterogeneity in human colorectal tumors. Nature Publishing Group, 49(5):708–718, May 2017.

[52] Qin Li, Anders Wennborg, Erik Aurell, Erez Dekel, Jie-Zhi Zou, Yuting Xu, Sui Huang, and Ingemar Ernberg. Dynamics inside the cancer cell attractor reveal cell heterogeneity, limits of stability, and escape. Proc Natl Acad Sci USA, 113(10):2672–2677, March 2016.

[53] Jing Shan Lim, Alvaro Ibaseta, Marcus M Fischer, Belinda Cancilla, Gilbert O’Young, Sandra Cristea, Vincent C Luca, Dian Yang, Nadine S Jahchan, Cécile Hamard, Martine Antoine, Marie Wislez, Christina Kong, Jennifer Cain, Yu-Wang Liu, Ann M Kapoun, K Christopher Garcia, Timothy Hoey, Christopher L Murriel, and Julien Sage. Intratumoural heterogeneity generated by Notch signalling promotes small-cell lung cancer. Nature, 545(7654):360–364, May 2017.

[54] Iain C Macaulay, Valentine Svensson, Charlotte Labalette, Lauren Ferreira, Fiona Hamey, Thierry Voet, Sarah A Teichmann, and Ana Cvejic. Single-Cell RNA-Sequencing Reveals a Continuous Spectrum of Differentiation in Hematopoietic Cells. Cell Rep, 14(4):1–13, January 2016.

[55] M C Mackey. Unified hypothesis for the origin of aplastic anemia and periodic hematopoiesis. Blood, 51(5):941–956, May 1978.

[56] M C Mackey. Cell kinetic status of haematopoietic stem cells. Cell Prolif, 34(2):71–83, April 2001.

[57] Nemanja D Marjanovic, Robert A Weinberg, and Christine L Chaffer. Cell plasticity and heterogeneity in cancer. Clin Chem, 59(1):168–179, December 2012.

[58] Joan Massague. TGF*β* signalling in context. Nat Rev Mol Cell Biol, 13(10):616–630, October 2012.

[59] Nicholas McGranahan and Charles Swanton. Clonal Heterogeneity and Tumor Evolution: Past, Present, and the Future. Cell, 168(4):613–628, February 2017.

[60] Alexander Meissner. Epigenetic modifications in pluripotent and differentiated cells. Nat Biotechnol, 28(10):1079–1088, October 2010.

[61] Luis G LG Morelli, Koichiro K Uriu, Saúl S Ares, and Andrew C AC Oates. Computational approaches to developmental patterning. Science, 336(6078):187–191, April 2012.

[62] R Morris, I Sancho-Martinez, T O Sharpee, and J C Izpisua Belmonte. Mathematical approaches to modeling development and reprogramming. Proc Natl Acad Sci USA, 111(14):5076–5082, April 2014.

[63] A Nakao, M Afrakhte, A Moren, T Nakayama, J L Christian, R Heuchel, S Itoh, N Kawabata, N E Heldin, C H Heldin, and P tenDijke. Identification of Smad7, a TGF beta-inducible antagonist of TGF-beta signalling. Nature, 389(6651):631–635, 1997.

[64] Nicholas Navin, Jude Kendall, Jennifer Troge, Peter Andrews, Linda Rodgers, Jeanne McIndoo, Kerry Cook, Asya Stepansky, Dan Levy, Diane Esposito, Lakshmi Muthuswamy, Alex Krasnitz, W Richard McCombie, James Hicks, and Michael Wigler. Tumour evolution inferred by single-cell sequencing. Nature, 472(7341):90–94, April 2011.

[65] Sonia Nestorowa, Fiona K Hamey, Blanca Pijuan Sala, Evangelia Diamanti, Mairi Shepherd, Elisa Laurenti, Nicola K Wilson, David G Kent, and Berthold Göttgens. A single-cell resolution map of mouse hematopoietic stem and progenitor cell differentiation. Blood, 128(8):e20–31, August 2016.

[66] Faiyaz Notta, Sasan Zandi, Naoya Takayama, Stephanie Dobson, Olga I Gan, Gavin Wilson, Kerstin B Kaufmann, Jessica McLeod, Elisa Laurenti, Cyrille F Dunant, John D McPherson, Lincoln D Stein, Yigal Dror, and John E Dick. Distinct routes of lineage development reshape the human blood hierarchy across ontogeny. Science, 351(6269):aab2116–aab2116, January 2016.

[67] David M Ornitz and Nobuyuki Itoh. Fibroblast growth factors. Genome Biol., 2(3):reviews 3005.1–3005.12, 2001.

[68] David-Emlyn Parfitt and Magdalena Zernicka-Goetz. Epigenetic Modification Affecting Expression of Cell Polarity and Cell Fate Genes to Regulate Lineage Specification in the Early Mouse Embryo. Mol Biol Cell, 21(15):2649–2660, 2010.

[69] Mireya Plass, Jordi Solana, F Alexander Wolf, Salah Ayoub, Aristotelis Misios, Petar Glažar, Benedikt Obermayer, Fabian J Theis, Christine Kocks, and Nikolaus Rajewsky. Cell type atlas and lineage tree of a whole complex animal by single-cell transcriptomics. Science, 360(6391):eaaq1723–12, May 2018.

[70] Anna Portela and Manel Esteller. Epigenetic modifications and human disease. Nat Biotechnol, 28(10):1057–1068, September 2010.

[71] Aline V Probst, Elaine Dunleavy, and Geneviéve Almouzni. Epigenetic inheritance during the cell cycle. Nat Rev Mol Cell Biol, 10(3):192–206, February 2009.

[72] A Rudenko and L H Tsai. Epigenetic regulation in memory and cognitive disorders. Neuroscience, 264:51–63, April 2014.

[73] Anguraj Sadanandam, Costas A Lyssiotis, Krisztian Homicsko, Eric A Collisson, William J Gibb, Stephan Wullschleger, Liliane C Gonzalez Ostos, William A Lannon, Carsten Grotzinger, Maguy Del Rio, Benoit Lhermitte, Adam B Olshen, Bertram Wiedenmann, Lewis C Cantley, Joe W Gray, and Douglas Hanahan. A colorectal cancer classification system that associates cellular phenotype and responses to therapy. Nat Med, 19(5):619–625, May 2013.

[74] Francisco Sanchez-Vega, Marco Mina, Joshua Armenia, Konnor C La, Sofia Dimitriadoy, David L Liu, Havish S Kantheti, Sadegh Saghafinia, Foysal Da- ian, Qingsong Gao, Matthew H Bailey, Wen-Wei Liang, Steven M Foltz, Ilya Shmulevich, Li Ding, Zachary Heins, Benjamin Gross, Hongxin Zhang, Ritika Kundra, Istemi Bahceci, Leonard Dervishi, Ugur Dogrusoz, Wanding Zhou, Gregory P Way, Casey S Greene, Yonghong Xiao, Chen Wang, Antonio Iavarone, Alice H Berger, Trever G Bivona, Alexander J Lazar, Gary D Hammer, Thomas Giordano, Lawrence N Kwong, Grant McArthur, Chenfei Huang, Aaron D Tward, Mitchell J Frederick, Frank McCormick, The Cancer Genome Atlas Research Network, Samantha J Caesar-Johnson, John A Demchok, Ina Felau, Melpomeni Kasapi, Martin L Ferguson, Carolyn M Hutter, Heidi J Sofia, Roy Tarnuzzer, Zhining Wang, Liming Yang, Jean C Zenklusen, Jiashan Julia Zhang, Sudha Chudamani, Jia Liu, Laxmi Lolla, Rashi Naresh, Todd Pihl, Qiang Sun, Yunhu Wan, Ye Wu, Juok Cho, Timothy DeFreitas, Scott Frazer, Nils Gehlenborg, Gad Getz, David I Heiman, Jaegil Kim, Michael S Lawrence, Pei Lin, Sam Meier, Michael S Noble, Gordon Saksena, Doug Voet, Hailei Zhang, Brady Bernard, Nyasha Chambwe, Varsha Dhankani, Theo Knijnenburg, Roger Kramer, Kalle Leinonen, Yuexin Liu, Michael Miller, Sheila Reynolds, Vesteinn Thorsson, Wei Zhang, Rehan Akbani, Bradley M Broom, Apurva M Hegde, Zhenlin Ju, Rupa S Kanchi, Anil Korkut, Jun Li, Han Liang, Shiyun Ling, Wenbin Liu, Yiling Lu, Gordon B Mills, Kwok-Shing Ng, Arvind Rao, Michael Ryan, Jing Wang, John N Weinstein, Jiexin Zhang, Adam Abeshouse, Debyani Chakravarty, Walid K Chatila, Ino de Bruijn, Jianjiong Gao, Benjamin E Gross, Zachary J Heins, Konnor La, Marc Ladanyi, Augustin Luna, Moriah G Nissan, Angelica Ochoa, Sarah M Phillips, Ed Reznik, Chris Sander, Robert Sheridan, S Onur Sumer, Yichao Sun, Barry S Taylor, Jioajiao Wang, Pavana Anur, Myron Peto, Paul Spellman, Christopher Benz, Joshua M Stuart, Christopher K Wong, Christina Yau, D Neil Hayes, Joel S Parker, Matthew D Wilkerson, Adrian Ally, Miruna Balasundaram, Reanne Bowlby, Denise Brooks, Rebecca Carlsen, Eric Chuah, Noreen Dhalla, Robert Holt, Steven J M Jones, Katayoon Kasaian, Darlene Lee, Yussanne Ma, Marco A Marra, Michael Mayo, Richard A Moore, Andrew J Mungall, Karen Mungall, A Gordon Robertson, Sara Sadeghi, Jacqueline E Schein, Payal Sipahimalani, Angela Tam, Nina Thiessen, Kane Tse, Tina Wong, Ashton C Berger, Rameen Beroukhim, Andrew D Cherniack, Carrie Cibulskis, Stacey B Gabriel, Galen F Gao, Gavin Ha, Matthew Meyerson, Steven E Schumacher, Juliann Shih, Melanie H Kucherlapati, Raju S Kucherlapati, Stephen Baylin, Leslie Cope, Ludmila Danilova, Moiz S Bootwalla, Phillip H Lai, Dennis T Maglinte, David J Van Den Berg, Daniel J Weisenberger, J Todd Auman, Saianand Balu, Tom Bodenheimer, Cheng Fan, Katherine A Hoadley, Alan P Hoyle, Stuart R Jefferys, Corbin D Jones, Shaowu Meng, Piotr A Mieczkowski, Lisle E Mose, Amy H Perou, Charles M Perou, Jeffrey Roach, Yan Shi, Janae V Simons, Tara Skelly, Matthew G Soloway, Donghui Tan, Umadevi Veluvolu, Huihui Fan, Toshinori Hinoue, Peter W Laird, Hui Shen, Wanding Zhou, Michelle Bellair, Kyle Chang, Kyle Covington, Chad J Creighton, Huyen Dinh, HarshaVardhan Doddapaneni, Lawrence A Donehower, Jennifer Drummond, Richard A Gibbs, Robert Glenn, Walker Hale, Yi Han, Jianhong Hu, Viktoriya Korchina, Sandra Lee, Lora Lewis, Wei Li, Xiuping Liu, Margaret Morgan, Donna Morton, Donna Muzny, Jireh Santibanez, Margi Sheth, Eve Shinbrot, Linghua Wang, Min Wang, David A Wheeler, Liu Xi, Fengmei Zhao, Julian Hess, Elizabeth L Appelbaum, Matthew Bailey, Matthew G Cordes, Catrina C Fronick, Lucinda A Fulton, Robert S Fulton, Cyriac Kandoth, Elaine R Mardis, Michael D McLellan, and Christopher… Miller. Oncogenic Signaling Pathways in The Cancer Genome Atlas. Cell, 173(2):321–337.e10, April 2018.

[75] António J M Santos, Yuan-Hung Lo, Amanda T Mah, and Calvin J Kuo. The Intestinal Stem Cell Niche: Homeostasis and Adaptations. Trends Cell Biol, 28(12):1062–1078, December 2018.

[76] Albert Serra-Cardona and Zhiguo Zhang. Replication-Coupled Nucleosome Assembly in the Passage of Epigenetic Information and Cell Identity. Trends Biochem Sci, 43(2):136–148, February 2018.

[77] Jeffrey A Simon and Carol A Lange. Roles of the EZH2 histone methyltransferase in cancer epigenetics. Mutat Res, 647(1-2):21–29, November 2008.

[78] Alexander P Sobinoff and Hilda A Pickett. Alternative Lengthening of Telomeres: DNA Repair Pathways Converge. Trends in Genetics, 33(12):921–932, December 2017.

[79] You Song, Honglei Ren, and Jinzhi Lei. Collaborations between CpG sites in DNA methylation. Int. J. Mod. Phys. B, 31(2):1750243, August 2017.

[80] Rama Soundararajan, Anurag N Paranjape, Sankar Maity, Ana Aparicio, and Sendurai A Mani. EMT, stemness and tumor plasticity in aggressive variant neuroendocrine prostate cancers. Bba-Rev Cancer, 1870(2):229–238, December 2018.

[81] Mario L Suvà, Nicolo Riggi, and Bradley E Bernstein. Epigenetic reprogramming in cancer. Science, 339(6127):1567–1570, March 2013.

[82] Wai Leong Tam and Robert A Weinberg. The epigenetics of epithelial-mesenchymal plasticity in cancer. Nat Med, 19(11):1438–1449, 2013.

[83] Michael J Topper, Michelle Vaz, Katherine B Chiappinelli, Christina E DeStefano Shields, Noushin Niknafs, Ray-Whay Chiu Yen, Alyssa Wenzel, Jessica Hicks, Matthew Ballew, Meredith Stone, Phuoc T Tran, Cynthia A Zahnow, Matthew D Hellmann, Valsamo Anagnostou, Pamela L Strissel, Reiner Strick, Victor E Velculescu, and Stephen B Baylin. Epigenetic Therapy Ties MYC Depletion to Reversing Immune Evasion and Treating Lung Cancer. Cell, 171(6):1284–1300.e21, November 2017.

[84] Maria-Elena Torres-Padilla, David-Emlyn Parfitt, Tony Kouzarides, and Magdalena Zernicka-Goetz. Histone arginine methylation regulates pluripotency in the early mouse embryo. Nature, 445(7124):214–218, January 2007.

[85] Lars Velten, Simon F Haas, Simon Raffel, Sandra Blaszkiewicz, Saiful Islam, Bianca P Hennig, Christoph Hirche, Christoph Lutz, Eike C Buss, Daniel Nowak, Tobias Boch, Wolf-Karsten Hofmann, Anthony D Ho, Wolfgang Huber, Andreas Trumpp, Marieke A G Essers, and Lars M Steinmetz. Human haematopoietic stem cell lineage commitment is a continuous process. Nat Cell Biol, 19(4):271–281, March 2017.

[86] Robert A Weinberg. Using maths to tackle cancer. Nature, 449(7):978–981, October 2007.

[87] Benjamin Werner, David Dingli, Tom Lenaerts, Jorge M Pacheco, and Arne Traulsen. Dynamics of Mutant Cells in Hierarchical Organized Tissues. PLoS Comput Biol, 7(12):e1002290–9, December 2011.

[88] Benjamin Werner, Jacob G Scott, Andrea Sottoriva, Alexander R A Anderson, Arne Traulsen, and Philipp M Altrock. The Cancer Stem Cell Fraction in Hierarchically Organized Tumors Can Be Estimated Using Mathematical Modeling and Patient-Specific Treatment Trajectories. Cancer Research, 76(7):1705–1713, March 2016.

[89] Wendy Weston, Jennifer Zayas, Ruben Perez, John George, and Roland Jurecic. Dynamic equilibrium of heterogeneous and interconvertible multipotent hematopoietic cell subsets. Sci Rep, 4:5199–5199, June 2014.

[90] H Wu, X-Y Zhang, Z Hu, Q Hou, H Zhang, Y Li, S Li, J Yue, Z Jiang, S M Weissman, X Pan, B-G Ju, and S Wu. Evolution and heterogeneity of non-hereditary colorectal cancer revealed by single-cell exome sequencing. Oncogene, 36(20):2857–2867, May 2017.

[91] Hao Wu and Yi Zhang. Reversing DNA methylation: mechanisms, genomics, and biological functions. Cell, 156(1-2):45–68, January 2014.

[92] Kan Xie, Devon P Ryan, Brandon L Pearson, Kristin S Henzel, Frauke Neff, Ramon O Vidal, Magali Hennion, Isabelle Lehmann, Melvin Schleif, Susanne Schröder, Thure Adler, Birgit Rathkolb, Jan Rozman, Anna-Lena Schütz, Cornelia Prehn, Michel E Mickael, Marco Weiergräber, Jerzy Adamski, Dirk H Busch, Gerhard Ehninger, Anna Matynia, Walker S Jackson, Eckhard Wolf, Helmut Fuchs, Valerie Gailus-Durner, Stefan Bonn, Martin Hrabě de Angelis, and Dan Ehninger. Epigenetic alterations in longevity regulators, reduced life span, and exacerbated aging-related pathology in old father offspring mice. Proc Natl Acad Sci USA, 115(10):E2348–E2357, March 2018.

[93] Boris Zhivotovsky and Guido Kroemer. Apoptosis and genomic instability. Nat Rev Mol Cell Biol, 5(9):752–762, September 2004.

